# Unique role and vulnerability of EP300 KIX domain in small-cell lung cancer

**DOI:** 10.1101/2021.06.03.446569

**Authors:** Kee-Beom Kim, Ashish Kabra, Dong-Wook Kim, Yongming Xue, Pei-Chi Hou, Yunpeng Zhou, Leilani Miranda, Xiaobing Shi, Timothy P. Bender, John H. Bushweller, Kwon-Sik Park

## Abstract

EP300 (E1A binding protein p300) is a versatile transcription co-activator important in cell proliferation and differentiation. The gene *EP300* is frequently mutated in diverse cancer types, including small-cell lung cancer (SCLC). While it is widely believed that these mutations result in loss of EP300 function, the impact on SCLC pathogenesis remains largely unknown. Here we demonstrate that mutant EP300 variants lacking histone acetyltransferase (HAT) domain accelerate tumor development in autochthonous mouse models of SCLC. However, unexpectedly, complete knockout of *Ep300* suppresses tumor development and inhibits proliferation of both human and mouse SCLC cells. Genetic dissection of EP300 domains identifies kinase-inducible domain (KID)-interacting (KIX) domain, specifically its interaction with transcription factors such as CREB1 and MYB, as the determinant of pro-tumorigenic activity. Blockade of the KIX-mediated protein interactions using a small molecule and a recombinant peptide mimicking the KIX-binding sequences of EP300-interacting partners inhibits the growth of SCLC cells. These findings identify domain-specific roles of EP300 in SCLC and unique vulnerability of the EP300 KIX domain to potential therapeutics.

## Introduction

EP300 and CREBBP are closely related transcription coactivators important in cell proliferation and differentiation (Goodman and Smolik, 2000) and are frequent targets of various oncogenic insults. Oncogenic viral proteins coopt coactivator functions of EP300/CREBBP to promote cellular transformation (Kasper et al., 2006; MacPherson et al., 2007), while loss-of-function mutations in *EP300/CREBBP* are thought to promote numerous cancer types including lymphoma and small-cell lung cancer (SCLC) (Augert et al., 2017; Cancer Genome Atlas Research, 2014; George et al., 2015; Horton et al., 2017; Jia et al., 2018; Jiang et al., 2017; Mullighan et al., 2011; Pasqualucci et al., 2011; Peifer et al., 2012; Rudin et al., 2012; Zhang et al., 2017). However, the mechanisms of EP300/CREBBP mutation-driven tumorigenesis, as well as the individual contributions of the two coactivators, remain poorly defined in light of their numerous binding partners and overlapping functions across cell types and biological contexts (Attar and Kurdistani, 2017). Here we show that mutant EP300 variants lacking histone acetyltransferase domain (HAT) domain accelerate tumor development in autochthonous mouse models of SCLC, whereas complete knockout of *Ep300* suppresses tumor development and inhibits proliferation of SCLC cells. We identify the kinase-inducible domain-interacting (KIX) domain of EP300, specifically its interaction with binding partners, as the source of pro-tumorigenic activity and demonstrate that it can be inhibited using both a small molecule and a recombinant peptide. Thus, by synthesizing the SCLC genome data with experimental data from cancer cells and genetically engineered mouse models, we identify a novel, domain-specific role of EP300 in SCLC and a potential approach to the development of targeted therapies.

## Results

### EP300 knockout suppresses SCLC

To begin to determine role of EP300 in SCLC development *in vivo*, we conditionally deleted *Ep300* in two strains of genetically engineered mouse models of SCLC, *Rb1*/*Trp53*/*Rbl2*-mutant (*RPR2*) and *Rb1*/*Trp53*/*Myc*-mutant (*RPM*) mice. In addition to deleting *Rb1* and *Trp53*, introduction of adenovirus Cre results in deletion of *Rbl2* in *RPR2* mice and induction of an oncogenic form of MYC (MYC^T58A^) in *RPM* mice, leading to development of lung tumors resembling the major molecular subtypes of human SCLC (Figure 1A and S1A) (Kasper et al., 2006; Schaffer et al., 2010; Sutherland et al., 2011). We performed intratracheal infection of Ad-CGRP-Cre that expresses Cre under the control of a neuroendocrine-specific Calcitonin-gene related peptide (CGRP) promoter (Mollaoglu et al., 2017), and 7 months later, observed development of nodular lung tumors in *Ep300^+/+^ RPR2* mice as expected; *Ep300^∆/∆^ RPR2* mice, however, developed only small lesions (Figures 1B and 1C). The small lesions and tumors from mice of both genotypes consisted of small cells with scanty cytoplasm that stained for CGRP, indicative of SCLC (Figure S1B). Likewise, 9 weeks after infection of Ad-CGRP-Cre, mice with *Ep300* deletion (*Ep300^∆/∆^ RPM*) developed fewer and smaller lesions in the lung than mice with intact EP300 (*Ep300^+/+^ RPM*) (Figure 1B, 1C, and S1C). Lesions in *Ep300^∆/∆^ RPR2* and *Ep300^∆/∆^ RPM* mice contained a smaller proportion of cells stained positive for a proliferation marker, phosphorylated histone H3 (pHH3), than tumors in *Ep300^+/+^ RPR2* and *Ep300^+/+^ RPM* mice (Figure S1D). Consistently, CRISPR-mediated knockout of EP300 in tumor cells derived from *RP (Rb1/Trp53-*mutant) and *RPR2* mice and in human SCLC lines significantly inhibited cell proliferation compared with control cells expressing a non-targeting gRNA (Figures 1D and S2A-D). Notably, knockout of *Crebbp*, the structural and functional homolog of *Ep300*, did not have a significant effect, and neither *Ep300* nor *Crebbp* knockout significantly affected the proliferation of non-small cell lung cancer cells (NSCLC, A549 and H1299) and human bronchial epithelial cells (HBEC) (Figures 1D and S2E). These findings indicate a unique requirement for EP300 in SCLC tumor development and cell proliferation, and suggest that EP300 and CREBBP may have non-overlapping roles.

**Figure 1.**
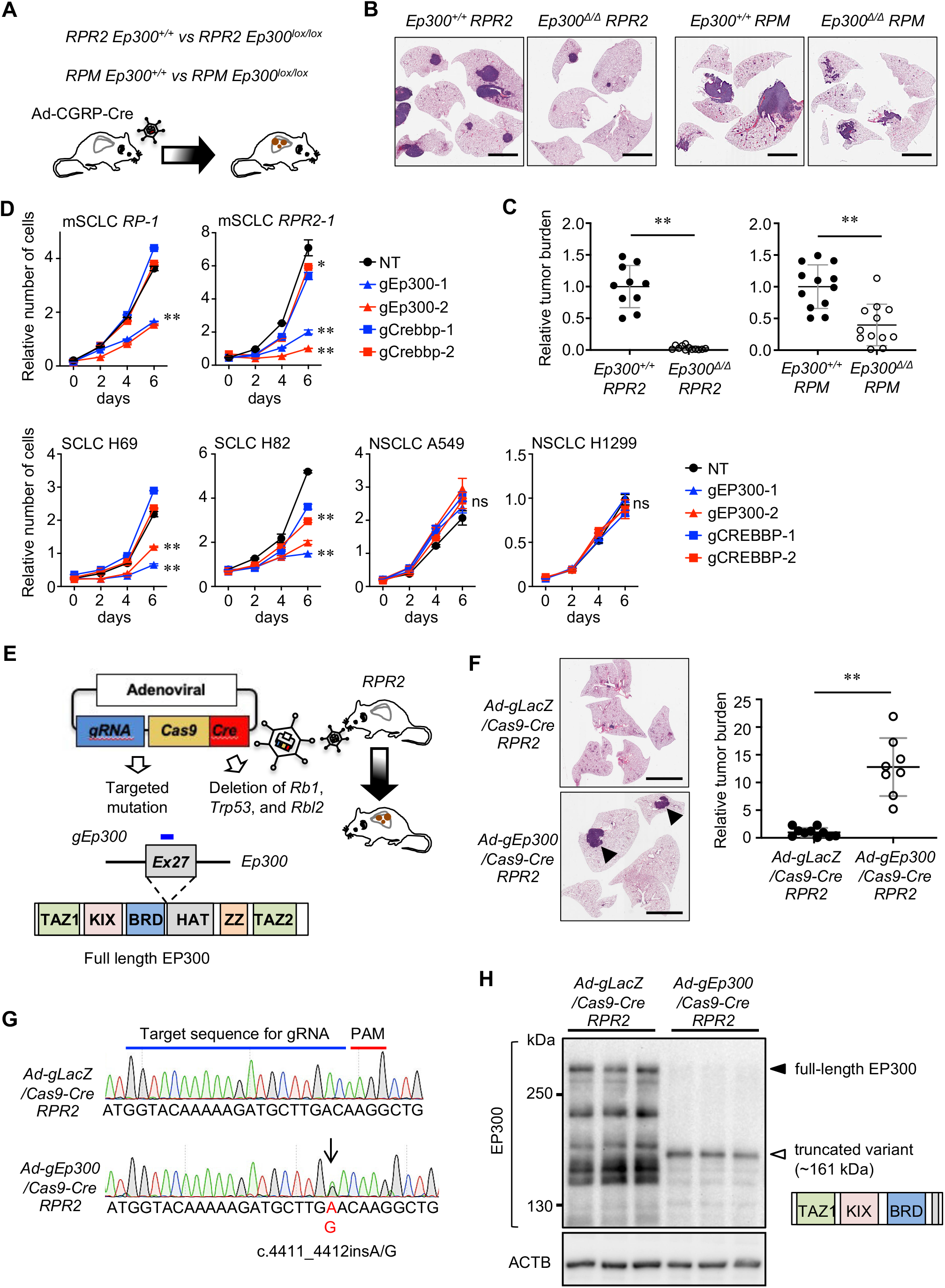
Tumor-suppressive effect of *Ep300* knockout and tumor-promoting effect of inactivating EP300 HAT domain on tumor development. **A,** Schematics of Cre-driven lung tumor development in *RPR*2 and *RPM* mice. **B,** Representative images of H&E-stained lung sections of *RPR*2 and *RPM* mice. **C,** Quantification of tumor burden (tumor area/lung area) in *Ep300^+/+^ vs. Ep300^∆/∆^ RPR2* (n=10 and n=12, respectively) and *Ep300^+/+^ vs. Ep300^∆/∆^ RPM* mice (n=12 tumors per genotype). **D,** Results of CellTiter-Glo assays for mouse SCLC cells (RP-1 and RPR2-1), derived from the tumors developed in *RP* (*Rb1/Trp53-* mutant) and *RPR2* mice, and human SCLC and NSCLC lines expressing CRISPR/Cas9 and gRNAs against *Ep300/EP300* or *Crebbp/CREBBP* or a non-targeting gRNA (NT). **E,** Schematics of adenoviral CRISPR/Cre hybrid vector and targeting of *Ep300* exon 27 in the lungs of *RPR2* mice. **F,** Representative images of H&E-stained lung sections of *RPR2* mice infected with Ad-gLacZ/Cas9-Cre *vs.* Ad-gEp300/Cas9-Cre virus expressing Cre and gRNAs against *LacZ* (gLacZ) and *Ep300* (gEp300) (left) and quantification of tumor burden (tumor area/lung area, n=10 and n=8, respectively) (right). Arrowheads indicate nodular lung tumors. **G,** Chromatograms showing the gRNA target sequences (chromosome 15: 81,525,569–81,525,602) in mice infected with Ad-gLacZ/Cas9-Cre or Ad-gEp300/Cas9-Cre. Arrow indicates the insertion mutation. **H,** Immunoblot for EP300 in primary cells derived from the lung tumors. Closed and open arrowheads indicate wild-type and variant EP300, respectively. ACTB blot for a protein loading control. *, p<0.05; **, p<0.01. Statistical tests performed using unpaired t-test (ns: not significant). Error bars represent standard deviation. Scale bars: B, F, 5mm.

### Inactivation of EP300 HAT promotes SCLC

The observation that *Ep300* knockout inhibits SCLC development in mice was unexpected and intriguing, given that knockout of its paralog *Crebbp* promotes tumor development (Jia et al., 2018). EP300 is expressed in SCLC patient tumors, where it frequently harbors mutations in the HAT domain (George et al., 2015). We thus postulated that introducing similar mutations in *Ep300* in mice would further promote tumor development. We designed an adenovirus CRISPR-Cre hybrid vector (Ad-gEp300/Cas9-Cre) that expresses Cre recombinase deleting the floxed alleles of *Rb1*, *Trp53*, and *Rbl2*, as well as Cas9 and a guide RNA against *Ep300* exon 27 encoding part of the HAT domain (Figure 1E). Ad-gLacZ/Cas9-Cre was used as a control that expresses guide RNA against a bacterial gene *LacZ*. After 7 months, *RPR2* mice infected with Ad-gEp300/Cas9-Cre developed larger lung tumor with a higher proportion of pHH3-positive cells than small lesions in mice infected with Ad-gLacZ/Cas9-Cre (Figures 1F and S3A). Sanger sequencing of tumor cells derived from mice infected with Ad-gEp300/Cas9-Cre consistently revealed insertion mutations (c.4411_4412insA/G) in exon 27 of *Ep300* resulting in a premature STOP codon and ~161 kDa truncated EP300 variant (Figures 1G and 3B). Consistently, immunoblot with antibodies against N-terminus of EP300 demonstrated loss of full length EP300 (300 kDa) and expression of the truncated variant in primary tumor cells from mice infected with Ad-gEp300/Cas9-Cre (Figure 1H). These findings suggest that CRISPR-mediated targeting of the HAT domain decreases the expression of full-length EP300 while inducing the expression of EP300 variants lacking most of the HAT domain and others domains at the C-terminus. Given that loss of *Ep300* abolished tumor development, while elimination of the HAT domain was sufficient to drive tumor development, we surmised that the pro-tumorigenic role for EP300 in SCLC is due to the activity of N-terminal domains, TAZ (transcriptional adaptor zinc-binding), KIX, and bromodomain (BRD).

### EP300 KIX is required for SCLC growth

We next sought to determine the contribution of the TAZ1, KIX, and BRD domains to tumor development using a cell-based model of SCLC development (Kim et al., 2016). Similar to mouse models, CRISPR-mediated targeting of *Ep300* exon 27 in *Rb1* and *Trp53*-deficient precancerous neuroendocrine cells (preSCs) results in loss of full-length EP300 and expression of EP300 variants lacking the HAT domain and the rest of C-terminus (Figures 2A, S4A, and S4B). Consistently, these *Ep300^∆HAT^*preSCs transformed into spheroids and highly tumorigenic cells (Figure 2B). We next performed CRISPR-mediated targeting of the exons encoding TAZ, KIX, BRD domains. Frame-shift mutations in exon 2, which result in deletion of all 3 domains, drastically reduced the ability of *Ep300^∆HAT^*preSCs to form colonies in soft agar compared with non-targeted control cells (Figure 2C, S4C, S4D, and S5A). A similar decrease in soft agar growth was seen with targeting of exon 9, expected to result in loss of function of the KIX and BRD (Figure 2C, S4C, S4D, and S5A). However, neither targeting of exon 16, which disrupts the function of BRD domain, nor chemical inhibition of BRD and HAT domains using selective inhibitors had any effect in these assays (Figure 2C, 2D, S4C, S4D, S5A, and S5B). These findings indicate that the KIX domain is necessary and sufficient for the oncogenic function of EP300 in SCLC. Notably, targeting the CREBBP KIX in *Ep300^∆HAT^*preSCs did not have any impact on soft agar growth (Figures 2E and S5C), further suggesting a specific role for the EP300 KIX in tumor development. Next we tested whether the role of the KIX domain is attributed to its interaction with transcription factors critical for cell proliferation and survival by individually knocking out the genes encoding known KIX-interacting factors (Figure 2F and S6A) (Chan and La Thangue, 2001). CRISPR-mediated targeting of CREB1, MYB, JUN, ATF1, and ATF4 inhibited the growth of *Ep300^∆HAT^*preSCs in soft agar, while targeting of SREBF2, RELA, GLI3, and KMT2A did not (Figures 2G and S6B). Notably, targeting the same factors in *Crebbp^KO^Ep300^∆HAT^*preSCs resulted in comparable results (Figures 2H and S6C). These results reveal a previously unidentified mechanism underlying SCLC development that is dependent on and unique to the EP300 KIX domain.

**Figure 2.**
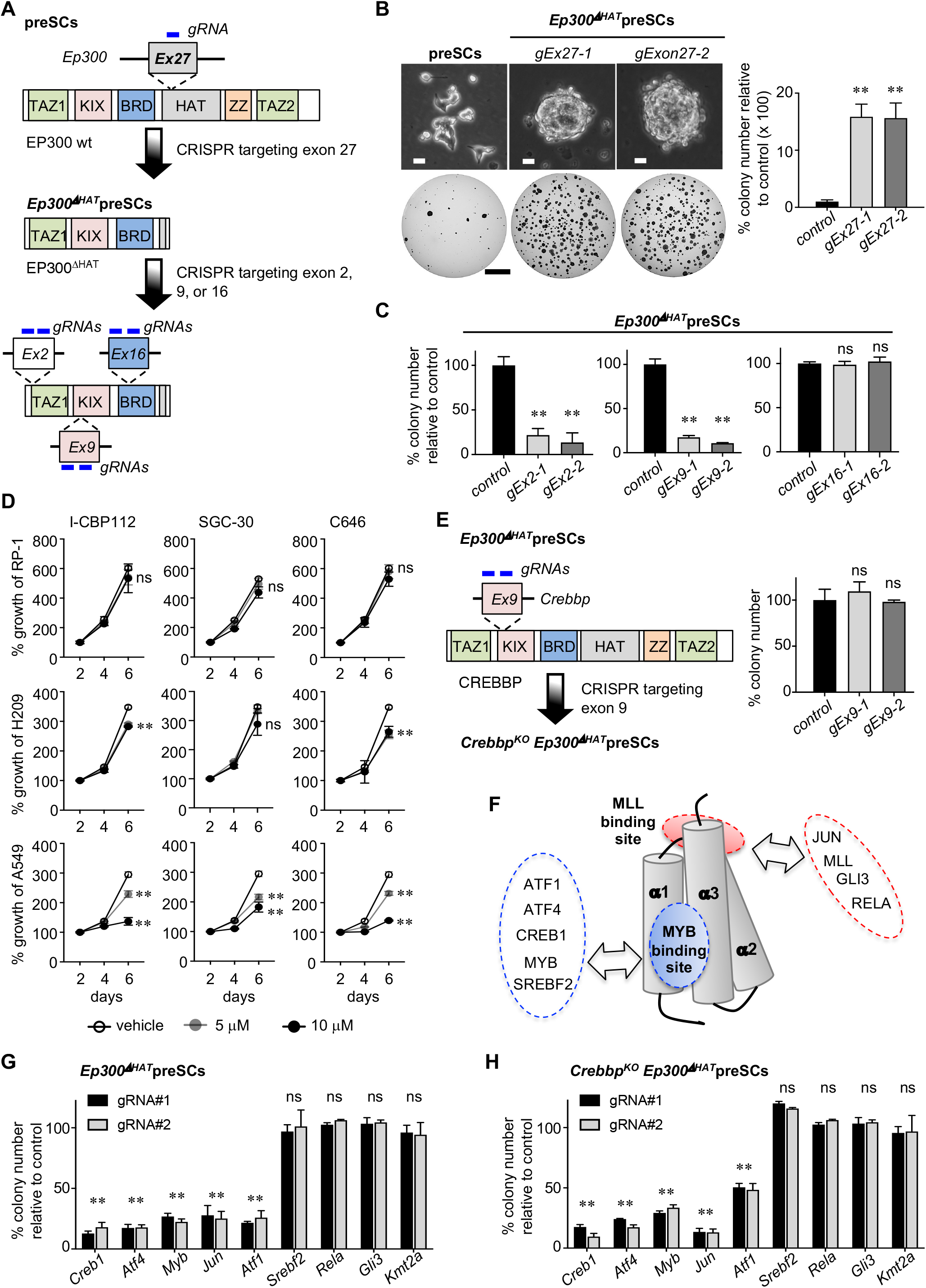
Role of EP300 KIX domain in SCLC cell proliferation. **A,** Schematics of generating *Ep300*^∆*HAT*^ preSCs by CRISPR targeting of *Ep300* exon 27, resulting in an EP300 variant lacking HAT and additional C-terminal domains (top), and CRISPR targeting of different exons encoding functional domains in *Ep300*^∆*HAT*^ preSCs. **B**, Representative images of control preSCs and *Ep300*^∆*HAT*^ preSCs in culture and soft agar (left) and quantification of colonies >0.2mm in diameter (right) (n=3 per cell type). **C,** Quantification of colonies (n=3 per cell type). **D,** Results of MTT viability assay of mouse SCLC cells and human SCLC and NSCLC cell lines treated with selective bromodomain inhibitors, I-CBP112 and SGC-CBP30, and with a selective inhibitor of HAT domain, C646, of CREBBP and EP300 (n=3 per cell type). **E,** Schematics of generating *Crebbp^KO^Ep300*^∆*HAT*^ preSCs (left) and quantification of soft agar colonies formed from *Crebbp* targeted and control *Ep300*^∆*HAT*^ cells (right) (n=3 per cell type). **F,** Cylinder representation of the structure of the KIX domain with MLL and MYB binding sites. Included in the dotted lines are the transcription factors known to bind to the MLL/MYB sites. **G, H,** Quantification of soft agar colonies formed from targeted and control *Ep300*^∆*HAT*^ preSCs and *Crebbp^KO^Ep300*^∆*HAT*^ preSCs. (n=3 per cell type). **, p<0.01. Statistical tests performed using unpaired t-test (ns: not significant). Error bars represent standard deviation. Scale bars; B, 10 µm (top), 5 mm (bottom).

### Disruption of KIX-mediated protein interaction inhibits SCLC growth

Transcription factors involved in proliferation and survival are known to bind to the KIX domain via common binding sites known as MLL and MYB sites (Figures 2F and 3A) (Chan and La Thangue, 2001). We thus sought to determine whether we could target these interactions to inhibit SCLC development and growth. We employed a fluorescence polarization assay (Scott et al., 2012) to measure the binding affinity of the KIX-mediated interactions using the purified KIX domains of EP300 and CREBBP and fluorophore-labeled KIX-binding sequences of MLL (MLL25) and MYB (MYB25). Purified EP300 KIX bound to MLL25 and MYB25 with higher affinity (K_d_ = 40 ± 14 μM and 3.8 ± 0.2 μM, respectively) than CREBBP KIX (K_d_ = 94 ± 11 μM and 22 ± 4.1 μM, respectively) (Figure 3B). These differential affinities have not been previously demonstrated but may result from differences in the amino acid sequences of the EP300 and CREBBP KIX domains. For example, in contrast to Asp647 in the CREBBP KIX domain, Ala647 in the EP300 KIX domain may mediate a hydrophobic interaction with Met303 of MYB25 that results in higher binding affinity between the EP300 KIX domain and its interacting partners (Figure 3A). Next, to test whether we could disrupt the interactions between the KIX domain and MLL and MYB, we generated a fusion peptide MLL28MYB30 (M/M) consisting of 28 residues from the MLL activation domain and 30 residues from the MYB activation domain linked by two glycine residues (GG) (Figure 3C). This design is based on data suggesting that MLL and MYB can bind cooperatively to the KIX domain to form a ternary complex that prevents other factors from binding to their respective sites in leukemia and other cell types (De Guzman et al., 2006) and that MYB and MLL could be tethered together via their interactions with Menin (Jin et al., 2010). As a control, we also made a variant of M/M (M/M-5A) with 5 alanine substitutions of critical residues important for the binding of MLL and MYB to the KIX domains. M/M inhibited binding of the FITC-labeled MYB and Rhodamine-labeled MLL peptides to the EP300 KIX domain with an IC_50_ value of 8.5 ± 0.7 μM and 8.3 ± 0.5 μM, respectively (Figure 3D). M/M also inhibited the binding of MYB and MLL peptides to the CREBBP KIX domain but at higher IC_50_ values, 22 ± 2.3 μM and 23 ± 1.3 μM, respectively. Importantly, M/M-5A did not have any effect even at high concentrations (Figure 3D). Immunoprecipitation followed by immunoblot confirmed the binding of M/M to both EP300 and CREBBP and indicated that, consistent with the results of the *in vitro* binding assay (Figure 3B), M/M bound preferentially to EP300 than to CREBBP (Figure 3E). Given that M/M appears to specifically inhibit EP300 KIX-mediated protein interactions, we expressed it in SCLC lines using a doxycycline-inducible lentivirus vector to determine the functional impact of disrupting these interactions (Figure S7A). M/M inhibited the ability of SCLC cells to form colonies in soft agar and their short-term proliferation, whereas M/M-5A did not have any significant effects (Figures 3F, S7B, and S7C). Notably, both M/M and M/M-5A did not detectably affect the growth of lung adenocarcinoma cell lines (Figures 3F and S7), which may be due in part to the low levels or absence of KIX domain-interacting proteins (Figure 3G). These results not only support the importance of the EP300 KIX domain for the growth of SCLC cells, but also suggest selective binding of the EP300 KIX domain to key interaction partners involved in proliferation and survival.

**Figure 3.**
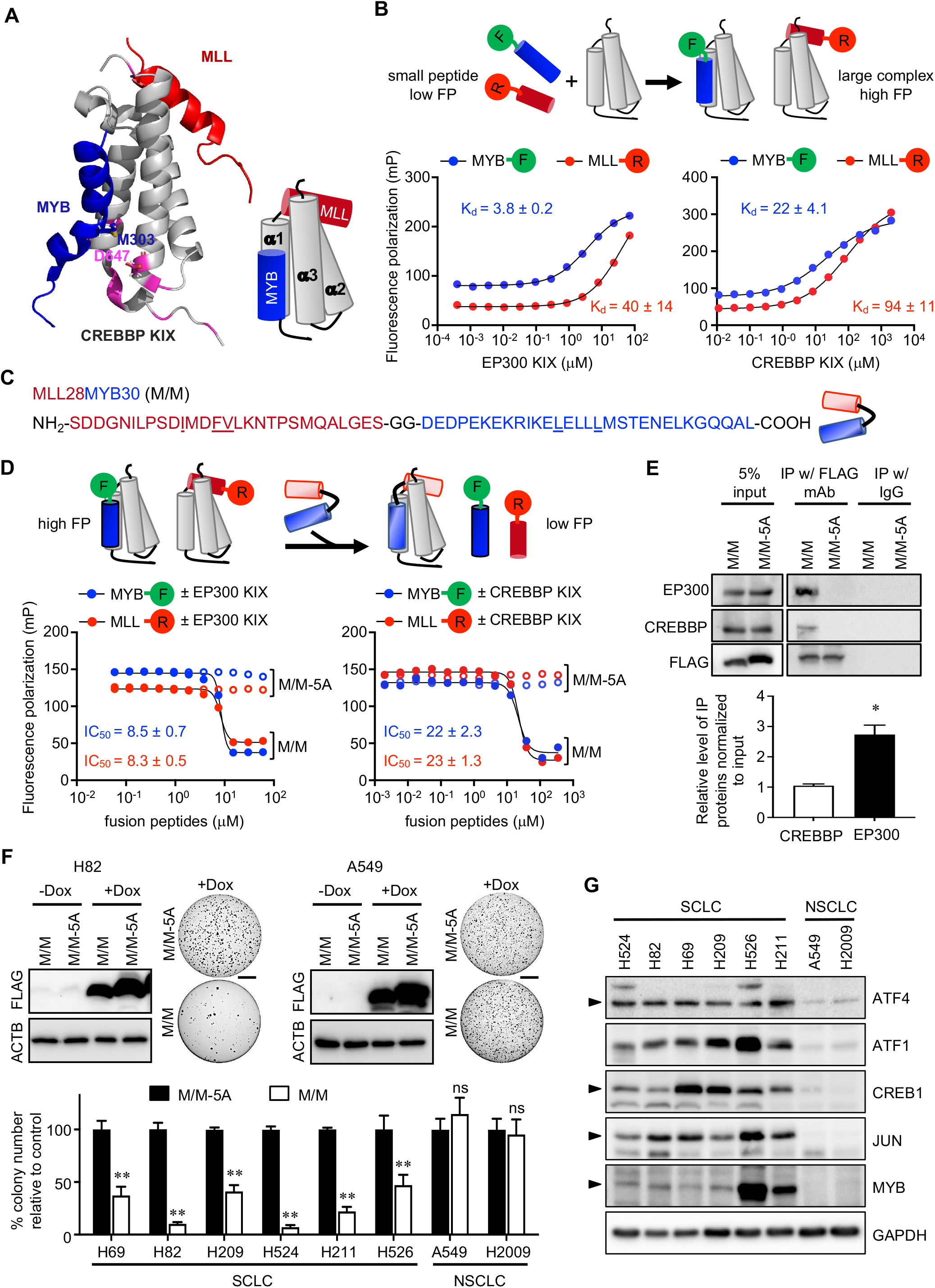
Effects of disrupting KIX domain-mediated protein interaction on cell proliferation. **A,** The structure of CREBBP KIX domain interacting with the MLL25 and MYB25 peptides representing 25 amino-acid fragments of MLL and MYB, respectively (PDB ID: 2AGH) (left) and cylinder diagram of the protein-peptide interactions (right). Residues highlighted in magenta represent key differences between the CREBBP and EP300 KIX domains. **B,** Schematic of the fluorescence polarization assay to examine binding with Rhodamine(R)-tagged MLL25 and FITC(F)-tagged MYB25 by the purified KIX domains of CREBBP and EP300 (top). K_d_ values indicate the binding affinities of the purified KIX domains for the fluorophore labeled peptides (bottom). **C,** Amino acid sequence of the fusion peptide MLL28MYB30 (M/M) with a GG linker (left) and diagram of M/M peptide with the linker (right). The underlined amino acids were replaced with alanine in the mutant variant M/M-5A. **D,** Schematic showing displacement of fluorophore labeled peptides by M/M (top). Inhibition curves (bottom) showing the impact of M/M and M/M-5A on binding of the EP300 and CREBBP KIX domains to fluorophore labeled peptides. **E,** Immunoblots for FLAG, EP300, and CREBBP in H524 cell extracts after immunoprecipitation (IP) with anti-FLAG antibodies or Isotype IgG. 5% of the cell extracts were used to show protein levels in input (top). Quantification of intensity of the bands following IP, relative to input band (n=3) (bottom). **F,** Immunoblots for FLAG in SCLC and NSCLC lines expressing FLAG-tagged M/M or M/M-5A under the control of doxycycline(dox)-inducible promoter, and images of soft agar colonies (top). Quantification of colonies >0.2mm in diameter (n=3) (bottom). **G,** Immunoblots for KIX domain-interacting proteins in SCLC and NSCLC lines. Arrowhead indicate specific bands. Statistical tests performed using unpaired t-test (ns: not significant). Error bars indicate the standard deviation in measurements. *, p<0.05; **, p<0.01. Scale bars: 5mm.

### KIX domain is vulnerable to chemical inhibition

In light of results from our proof-of-concept experiments, we next sought to explore the feasibility of using small molecules to block KIX domain-protein interactions and regulate cell growth. KG-501 (2-naphtol-AS-E-phosphate) and its derivative 666-15 have been shown to inhibit the interaction between CREB1 and the KIX domains of EP300 and CREBBP (Best et al., 2004; Xie et al., 2015). KG-501 inhibited the short-term growth of SCLC lines in culture and colony formation in soft agar, but did not have a significant effect on NSCLC cell lines (Figures 4A and S8A). Likewise, 666-15 inhibited the growth of SCLC lines but did not affect the growth of NSCLC cell lines (Figures 4B, S8B and S8C). Given its improved potency and bioavailability compared with KG-501 (Li et al., 2016; Xie et al., 2015), 666-15 was tested for its potential effect on tumor growth *in vivo*. Daily intraperitoneal treatment with 666-15 suppressed the formation of subcutaneous tumors derived from mouse SCLC cells in immune-competent mice (129S/B6 F1 hybrid), without significant weight loss (Figures 4C and S8D). Encouraged by this result, we then tested the effects on SCLC development of treating *RPR2* mice with daily 666-15, beginning 180 days after tumor induction (Figures 4D and 4E). A conditional luciferase allele *Rosa26*^lox-stop-lox-Luc^ carried by these mice allowed for bioluminescence imaging to monitor the tumorigenic growth of mutant cells. After 4 weeks of treatment, bioluminescence measurement indicated a lower tumor burden in the inhibitor-treated mice than the vehicle-treated controls (Figure 4D). Furthermore, a moderate but significant improvement in survival outcomes was observed in mice treated with 666-15 compared with the vehicle (Figure 4E). These results provide further evidence for the importance of the KIX domain for the growth of SCLC cells, and strongly support for the feasibility of targeting KIX-mediated interactions for the treatment of SCLC *in vivo*.

**Figure 4.**
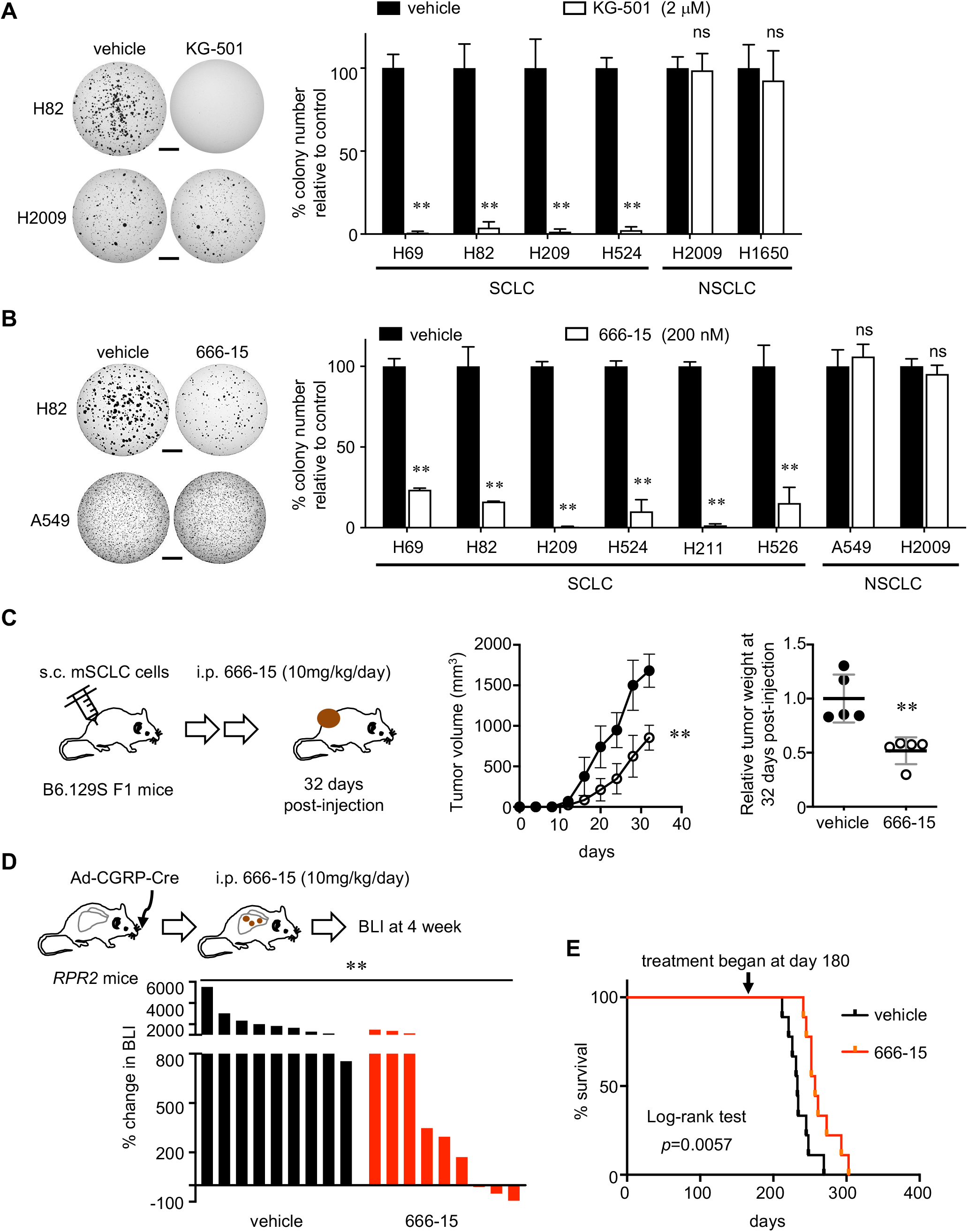
Chemical inhibition of the KIX domain suppresses SCLC cells proliferation and tumor development. **A, B,** Representative images of soft agar colonies (left) and quantification of colonies >0.2mm in diameter (n=3 per cell type) (right). Lung cancer cell lines were treated with 2 µM KG-501, 200 nM 666-15 in DMSO or equal volume DMSO (control) throughout the assay. This assay was repeated with comparable results. **C,** Schematic describing generation of allograft tumors in B6.129S F1 hybrid mice followed by treatment with 666-15 or vehicle control (left) and plots of tumor volumes (middle) and relative tumor weight (right) of allograft tumors formed in mice treated with vehicle or 666-15 (n=5 tumors per group). **D,** Schematic describing Cre-mediated induction of tumors in *RPR2* mice (top), quantification of changes in tumor volume per bioluminescence imaging (BLI), normalized to pre-treatment BLI levels (bottom). The vehicle or drug treatment began 180 days after Ad-CGRP-Cre infection for tumor induction. BLI was performed after 4 weeks of treatment. Each bar represents an individual tumor. **E,** Kaplan-Meier survival curves of *RPR2* mice treated with vehicle or 666-15 (n=9 tumors per genotype). Log-rank (Mantel-Cox) test was used to determine the significance of tumor-free survival between the cohorts. **, p<0.01. Statistical tests performed using unpaired t-test (ns: not significant). Error bar represents standard deviation. Scale bar: 5mm.

## Discussion

We report that EP300 plays both pro-tumorigenic and tumor-suppressive roles driven by the functions of KIX and HAT domains, respectively. This dual role of EP300 in SCLC is distinct from the tumor-suppressive role of CREBBP. This observation is consistent with the lack of mutations in *EP300* within the KIX domain coding sequence reported in SCLC patient tumors, while *CREBBP* harbors mutations along the entire length of coding sequence (Figure S9). In contrast, in lymphoma and other cancer types in which the functional loss of either EP300 or CREBBP is thought to promote tumorigenesis, the pattern of mutations in *EP300* and *CREBBP* is similar (Figure S9). The necessity of the EP300 KIX domain appears unique to SCLC in the context of cancer, as *Ep300* knockout did not affect development of *Kras*^G12D^-driven lung adenocarcinoma in mice (Figure S2F) and inhibition of both EP300 and CREBBP KIX domains did not affect proliferation of lung adenocarcinoma cells (Figures 3 and 4). Only the combined loss of EP300 and CREBBP, specifically the loss of BRD and HAT domain activities, suppressed the proliferation of *Kras*-mutant lung adenocarcinoma cells and diffuse large B cell lymphoma (Meyer et al., 2019; Ogiwara et al., 2016). Nonredundant functions of EP300 and CREBBP KIX domains has been documented in hematopoiesis for which the EP300 KIX domain, not the CREBBP KIX, is required (Kasper et al., 2002). We surmise that such unique dependency on EP300 KIX domain may be due in part to the ~6-fold higher affinity of the EP300 KIX for pro-tumorigenic transcription factors such as MYB, compared with the CREBBP KIX domain, and the cell-type specificity of EP300 dependency may be due in part to significantly higher levels of the KIX domain-interacting factors in SCLC than NSCLC cells (Figure 3). The impact of this difference in binding affinity on the interactions of EP300 and CREBBP with context-dependent partners, and the downstream mechanisms leading to tumorigenesis, will be intriguing to characterize in future studies. Further, the dependency of SCLC development on interactions mediated by the EP300 KIX domain represents a novel, actionable vulnerability in SCLC, a disease characterized predominantly by mutations in tumor suppressor genes and a lack of readily actionable oncogenic drivers.

## Acknowledgements

For the data of MSK-IMPACT Clinical Cohort Sequencing from cBioportal, we are grateful to the members of the Molecular Diagnostics Service in the Department of Pathology funded in part by the Marie-Josée and Henry R. Kravis Center for Molecular Oncology and the MSK Cancer Center Core Grant (NIH P30CA008748). We thank J. Hsu and H. Agaisse for reading the manuscript. We also thank C. Sonnet in Vector Development Laboratory at Baylor College of Medicine for the service of generating adenovirus CRISPR/Cre hybrid vector. This work was supported by NIH R01CA194461 and U01CA224293 (K.-S.P), UVA 3Cavalier (K.-S.P. and J.H.B), NIH R01GM100776 and R56AI108767 (T.P.B.), Adenoid Cystic Carcinoma Research Foundation (J.H.B.), NIH R01CA204020 (X.S.). We also thank the Research Histology Core at the UVA Cancer Center (NIH P30CA044579).

## Author contributions

K.-B.K., X.S., T.P.B., J.H.B. and K.-S.P. conceived the study and designed the experiments; K.-B.K., A.K., Y.X., D.-W.K., P.-C.H., Y.Z. and K.-S.P. performed biochemical and cell culture experiments. K.-B.K., L.M. and K.-S. P. performed animal experiments. A.K. and Y.Z. generated peptides and performed chemical and biochemical experiments. K.-B.K., X.S., T.P.B., J.H.B. and K.-S.P. data analysis and interpretation. K.-B.K., T.P.B., J.H.B. and K.-S.P. wrote the manuscript with help from all authors. All authors discussed the results and commented on the manuscript

## Methods

### Mouse strains, tumor induction, and allografts

Strains of *Trp53^lox^*, *Rb1^lox^*, *Rbl2^lox^*, *Ep300^lox^*, *H11^lox-stop-lox-MycT58A^*, and *Kras^LSL-G12D^* alleles were generously shared by Drs. Anton Berns, Tyler Jacks, Julien Sage, Paul Brindle, Trudy Oliver and Robert Wechsler-Reya, respectively. Compound transgenic mice *Rb1^lox/lox^ Trp53^lox/lox^ Rbl2^lox/lox^* (*RPR*) mice and *Rb1^lox/lox^ Trp53^lox/lox^ H11 ^lox-STOP-lox^* ^-*MycT58A*^ (RPM) mice have been previously described (Jackson et al., 2001; Kasper et al., 2006; MacPherson et al., 2007; Marino et al., 2000; Mollaoglu et al., 2017; Sage et al., 2003). For tumor induction, lungs of 10-week-old mice were infected with adenoviral Cre via intratracheal instillation as previously described and mice were aged 6 months (DuPage et al., 2009). Multiple cohorts of independent litters were analyzed to control for background effects, and both male and female mice were used. Ad-CGRP-Cre were purchased from the University of Iowa Gene Transfer Vector Core. Ad-gLacZ/Cas9-Cre and Ad-gEp300/Cas9-Cre particles were produced in Vector Development Laboratory at Baylor College of Medicine. For allograft experiments, 5.0 × 10^5^ murine cells were injected in the flanks of B6.129S F1 mice (Jackson Laboratory). Perpendicular tumor diameters were measured using calipers. Volume was calculated using the formula L × W^2^ × 0.52, where L is the longest dimension and W is the perpendicular dimension. The injected mice were maintained and observed for palpable tumors according to procedures approved by IACUC and euthanized when tumor size reached 1.5 cm in diameter, the endpoint of allograft study under the guideline of the institutional animal policy. The Kaplan–Meier curve was used to plot the time of survival (maximal tumor volume). For *in vivo* drug treatment experiments, *RPR* mice carrying *Rosa26^LSL-Luciferase^* were infected with Ad-CGRP-Cre and monitored by bioluminescence imaging to identify mice with tumor growth in the lungs. Mice anesthetized with isoflurane were injected with 150 mg/kg D-luciferin (firefly) potassium salt (GOLDBIO, LUCK), and bioluminescence was detected using IVIS Spectrum (PerkinElmer) over 20 minutes to record maximal radiance. The peak total flux values were assessed from the anatomical region of interest using Living Image 4.0 (PerkinElmer) and were used for analysis. 666-15 was dissolved in 1% N-methylpyrrolidone (NMP), 5% Tween-80 in H_2_O. The mice were i.p. injected with vehicle or 666-15 at 10mg/kg once a day for 7 days (Cancer Genome Atlas Research, 2014). Bioluminescence imaging was performed to follow tumor volume after treatment start, and weights were monitored weekly during the course of treatment.

### Histology and immunohistochemistry

Mouse tissues were fixed in 4% paraformaldehyde in phosphate-buffered saline before paraffin embedding. Five micrometer-thick tissue sections were stained with hematoxylin and eosin staining and immunostaining. For quantification of phosphorylated histone H3 (pHH3)-positive cells and CGRP staining, tumors of similar size and area were included. Macroscopic images of lung sections were acquired using Olympus MVX10. Images of H&E and immunostained tissues were acquired using Nikon Eclipse Ni-U microscope. Image analysis and automated quantification were performed using NIS-Elements Basic Research (Nikon)

### Plasmids and chemicals

pCW57.1, pL-CRISPR.EFS.tRFP, psPAX2 and pMD2.G plasmids were obtained from Addgene (#41393, #57819, #12259, #12260; the gifts from David Root, Benjami Ebert, Didier Trono). pCW-Cas9 plasmid was purchased from Addgene (#50661). LV-gRNA-zeocin plasmid was acquired from Mazhar Adli. DNA fragments corresponding to the coding region of MLL28MYB30 (M/M) and M/M-5A peptides were purchased from IDT, cloned in pENTR4 and subsequently in pCW57.1 using Gateway cloning kits (Thermo Fisher). Sequences of oligonucleotides for cloning and gRNAs are listed in the Supplementary Information. All sequences were verified by Sanger sequencing. BRD inhibitors (SGC-CBP30, I-CBP112) and HAT inhibitor (C646) were purchased from Sigma (SML1133, SML1134, 382113). Inhibitors of CREB1-EP300/CREBBP KIX domains, KG-501 and 666-15, were purchased from Sigma (70485, 538341).

### Cell culture, lentiviral infection, and transient transfection

Murine SCLC cells (mSCLC) and precancerous cells (preSC) were derived from lung tumors and early-stage neuroendocrine lesions, respectively, developed in the *Rb1/Trp53-*GEMM (Kim et al., 2016; Schaffer et al., 2010). Human cell lines, including H69, H82, H209, H524, A549, H526, H211, H2009, H1650 were the gifts from Adi Gazdar and John Minna (UT Southwestern), Julien Sage (Stanford), Hisashi Harada (Virginia Commonwealth University), and Christopher Vakoc (Cold Spring harbor). These cell lines were authenticated by profiling patterns of seventeen short tandem repeats (ATCC, 135-XV and 200FTA), and tested negative for *Mycoplasma* using PlasmoTest-Mycoplasma Detection Kit (InvivoGen, rep-pt1). Experiments were performed within the first few passages after thawing frozen cell stocks. Cells were cultured in RPMI1640 medium (Cellgro) supplemented with 10% bovine growth serum (GE Healthcare, SH30541.03) and 1% penicillin-streptomycin (Invitrogen, 15140-122). Puromycin (Thermo Fisher Scientific, A11138-03) or zeocin (Thermo Fisher Scientific, R25001) was used to select stably transduced cells following lentiviral infection. For lentivirus production, we transfected lentiviral plasmids with packaging plasmids in 293T cells using polyethylenimine (Sigma, 408727), harvested supernatants containing viral particles 48 and 72 hours later, and filtered through 0.45μm PVDF filter before adding to culture of target cells in the presence of 5μg/ml polybrene (Sigma, H9268). To generate KO cells, the parental cells were infected with LV-gRNA-zeocin vector carrying sgRNAs which target the sequence encoding each gene or empty vector (control) using Lipofectamine 2000 (Invitrogen, 52887) according to the manufacturer’s instructions. Forty-eight hours later, the positive cells were selected using zeocin. Sanger sequencing verified mutation in the target sequence in the genes and immunoblot validated loss of the proteins. For experiments in Figs 1c, d, transient transfection was performed using X-tremeGENE DNA transfection reagent (Roche) to generate KO cells. 48 h after transfection, cells were collected for Western blot analysis. For packaging lentivirus, 293T cells were co-transfected with pMD2.G, pPAX2 and LentiCRISPRv2 constructs using X-tremeGENE DNA transfection reagent (Roche). For infections, cells were incubated with viral supernatants in the presence of 8 mg/ml polybrene; after 48 hours, the infected cells were selected with blasticidin (10 μg/ml) or puromycin (2 μg/ml) for 4-6 days before experiments.

### Cell proliferation assays

Cells were seeded into a 96-well plate for 200~500 cells in 100 μL medium per well in triplicate for each time point and their proliferation was measured every one or two days depending on the growth rate of those cells. There is a tight linear relationship between cell number and the concentration of ATP measured in cell lysate. The bioluminescence-based reagents such as CellTiter-Glo (Promega) can detect ATP, which provides a sensitive readout of cell proliferation. To measure proliferation, 25 μL CellTiter-Glo was directly added into each well and the plate was placed on an orbit shaker for gentle rocking to induce cell lysis. Each plate was incubated at room temperature for 10 minutes to stabilize luminescent signal before it was read using Fluostar Omega plate reader. Alternatively, MTT assay was performed to measure relative cell viability using (3-[4,5-dimethylthiazol-2-yl]-2,5-diphenyltetrazolium bromide; thiazolyl blue) tetrazolium salt and plate reader. Cells were seeded at 1×10^4^ per well in 96-well plates at day 0. At times ranging from 0 to 4 days or 6 days post-seeding, MTT was added (10 μL, final concentration 0.5 mg/mL) and cells were then incubated for 4 hours at 37°C. OD measurements were determined with an ELISA reader at a wavelength of 590 nm. Background was accounted for by subtracting the value of a vehicle-containing blank from all values. For soft agar assay to measure clonal expansion, cells were seeded at 1X10^4^ cells per well in 0.5 mL of growth medium containing 0.35% low-melting-point agarose (Invitrogen, 16520-100) and seeded on top of a 0.5 mL base layer of medium containing 0.5% agar. The medium was regularly changed every 3 days during the experiments. Colonies were fixed with 10% MeOH, 10% acetic acid at room temperature for 10 minutes and stained with 1% MeOH, 1% Formaldehyde, and 0.05% crystal violet after. Low-melting-point agarose was premixed with RPMI 2X (Fisher Scientific, SLM202B) complemented with 20% FBS, 200 U/mL penicillin, and 200 μg/mL streptomycin. Cells were allowed to grow at 37°C with 5% CO2 for 3 weeks. Images of wells are acquired using Olympus MVX10 scope, and colonies from the whole field of image were counted using the imaging software NIS-Elements Basic Research (Nikon). All of the cell culture experiments were performed in triplicates and repeated for a minimum of two biological replicates. The cells were treated with doxycycline (Sigma, D9891) during the experiment.

### Immunoblot analysis and immunoprecipitation

Cells were lysed in RIPA buffer (50 mM Tris-HCl pH7.4, 150 mM NaCl, 2 mM EDTA, 1% NP-40, 0.1% SDS) and sonicated for 20s at 15% intensity. Generally, 15~30 μg protein lysate was loaded into various percentages of SDS-polyacrylamide gels for electrophoresis in PAGE running buffer (25 mM Tris, 192 mM glycine, 0.1% SDS, pH8.3). Then proteins were transferred to 0.2 or 0.45 μm PVDF membrane in transfer buffer (25 mM Tris, 192 mM glycine, 20% methanol). The membranes were blocked by TBST buffer (150 mM NaCl, 10 mM Tris pH8.0, 0.1% Tween20) supplemented with 5% non-fat milk for 1 hour at room temperature and then blotted with primary antibodies overnight at 4°C on an orbital shaker. After washing 5 times with TBST, membranes were incubated with secondary antibodies for 2 hours at room temperature and washed again with TBST 5 times. The membranes were then either used for ChemiDoc machine (Bio-Rad)-developing after incubating with ECL Western Blotting Detection Reagent (Thermo Fisher Scientific, 32106) for 5 minutes. For immunoprecipitation of protein complexes consisting of EP300/CREBBP-FLAG-tagged M/M fusion peptide, cells were lysed in RIPA buffer (50 mM Tris-HCl (pH 8.0), 150 mM NaCl, 0.1 % SDS, 0.5 % SDC, 1 % NP-40, 1X protease inhibitor cocktail, and 1 mM EDTA) and immunoprecipitated with anti-FLAG or anti-IgG antibodies in buffer (50 mM Tris-HCl (pH 7.5), 150 mM NaCl, 1 mM EDTA, 1 mM EGTA, 1 % Triton X-100, 1 mM PMSF, and 1X protease inhibitor cocktail) overnight at 4 °C. Protein A/G agarose beads (GenDEPOT, P9203) were then added for 4 h with agitation at 4 °C. Bound proteins were eluted and analyzed by immunoblotting with anti-FLAG, anti-EP300, and anti-CREBBP antibodies.

### Peptides and Fluorescence polarization assay

Human P300 KIX domain (87 residues spanning from 565-652) and the CBP KIX domain (87 residues spanning from 585-672) were expressed using a pMAL-c5E vector. MLL28MYB30 (wild-type and mutant) fusion peptides (28 residues from MLL activation domain and 30 residues from MYB activation domain linked by a GG linker) were expressed using a pET-32C vector (fused to thioredoxin). All proteins were expressed in *E. coli* Rosetta (DE3) cells and purified using affinity chromatography followed by anion-exchange chromatography. Fluorescence polarization (FP) assays were conducted in 96-well Costar black Polystyrene plates (Corning). N-terminally FITC labelled MYB (KEKRIKELELLLMSTENELKGQQAL) and Rhodamine labelled MLL peptides (SDDGNILPSDIMDFVLKNTPSMQAL) were used to detect KIX domain binding via changes in fluorescence polarization (FP). The FP assay mixture was composed of 50 mM HEPES pH 7.5, 50 mM NaCl, 12 μM and 45 μM purified P300 KIX domain or CBP KIX domain, respectively (~3 x K_d_ for KIX domain binding to the MYB peptide), 0.5 μM of each fluorescent peptide, and a variable concentration of purified peptide fusion inhibitor to make a total volume of 100 μL per well. After the addition of all assay components, the plates were incubated in the dark for the 30 minutes at room temperature. The plates were read using a PHERAstar plate reader (BMG Labtech). The FP values as a function of peptide fusion inhibitor concentration were fitted to a sigmoidal curve using Origin software.

### Statistical analysis

Statistical analyses and graphical presentation were performed with GraphPad Prism 8.2. The results are presented as the mean ± SD and evaluated using an unpaired Student t test (two-tailed; p<0.05 was considered to be significant). Kaplan–Meier curves of lung tumor-free survival were generated using Prism 8.2 (log-rank test; p<0.05 was considered to be significant).

---

**Figure S1.**
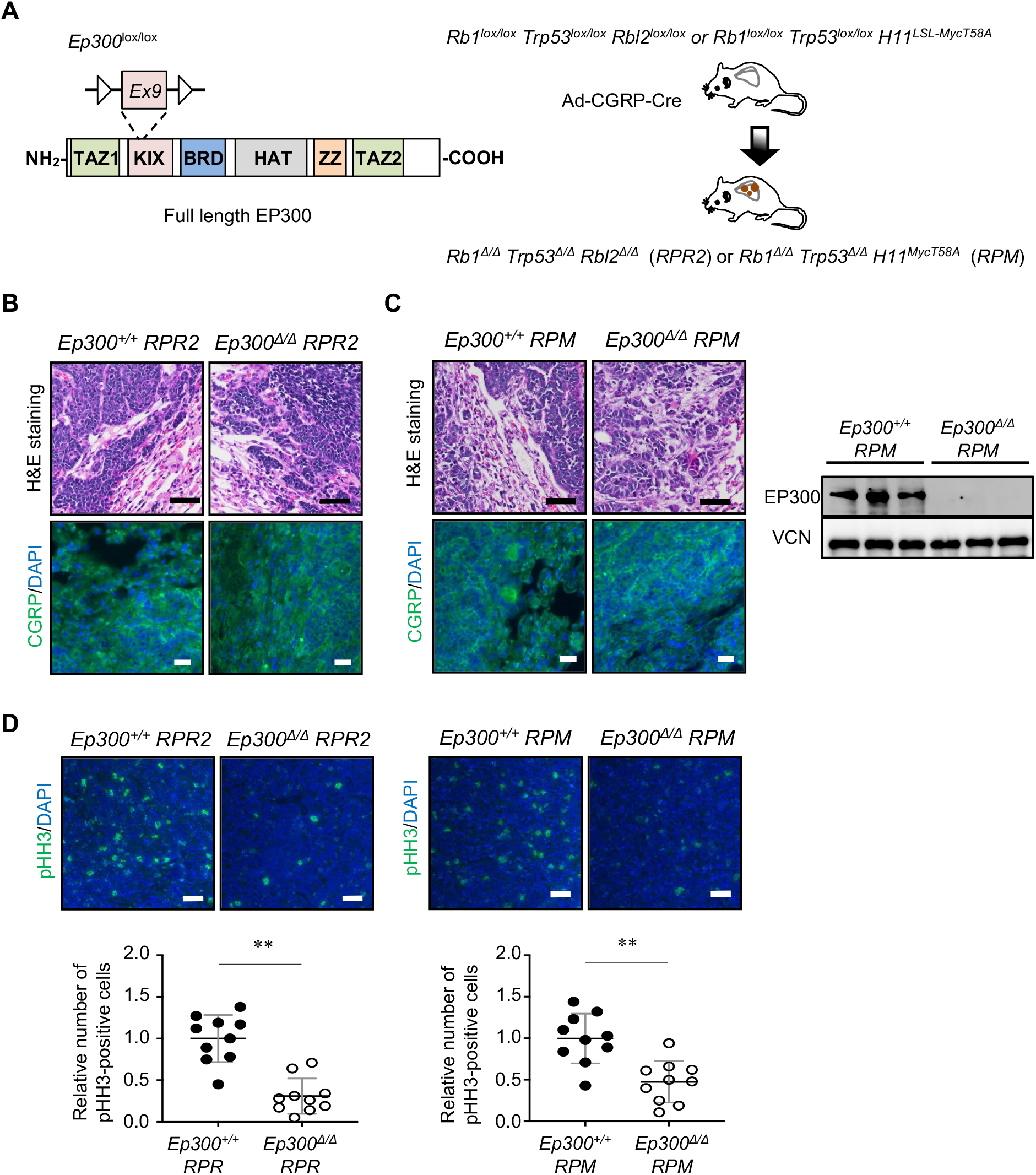
Autochthonous mouse models of SCLC with conditional deletion of EP300. **A,** Diagrams of the floxed *Ep300* allele and EP300 domains (left) and schematics of Ad-CGRP-Cre-driven tumor development in the lungs of *RPR*2 and *RPM* mice and (right). **B,** Representative images of H&E-stained lung sections of *Ep300*^+/+^ *vs. Ep300*^∆/∆^ *RPR2* mice (top) and immunostaining for CGRP (bottom). DAPI (blue) stains for nuclei. **C**, Representative images of H&E-stained lung sections of *Ep300*^+/+^ *vs. Ep300*^∆/∆^ *RPM* mice, and immunostaining for CGRP (left); immunoblot for EP300 in lung tumors developed in these mice (right). VCN (Vinculin) blot was used as a protein loading control. **D**, Representative images of immunostaining for phosphorylated histone H3 (pHH3) (top) and quantification of pHH3-positive cells per tumor area in *Ep300^+/+^ vs. Ep300^∆/∆^ RPR2* (n=10 tumors per genotype) and in *Ep300^+/+^ vs. Ep300^∆/∆^ RPM* mice (n=10 tumors per genotype) (bottom). **, p<0.01. Statistical tests performed using unpaired t-test. Error bars represent standard deviation. Scale bars: B-D, 50 mm.

**Figure S2.**
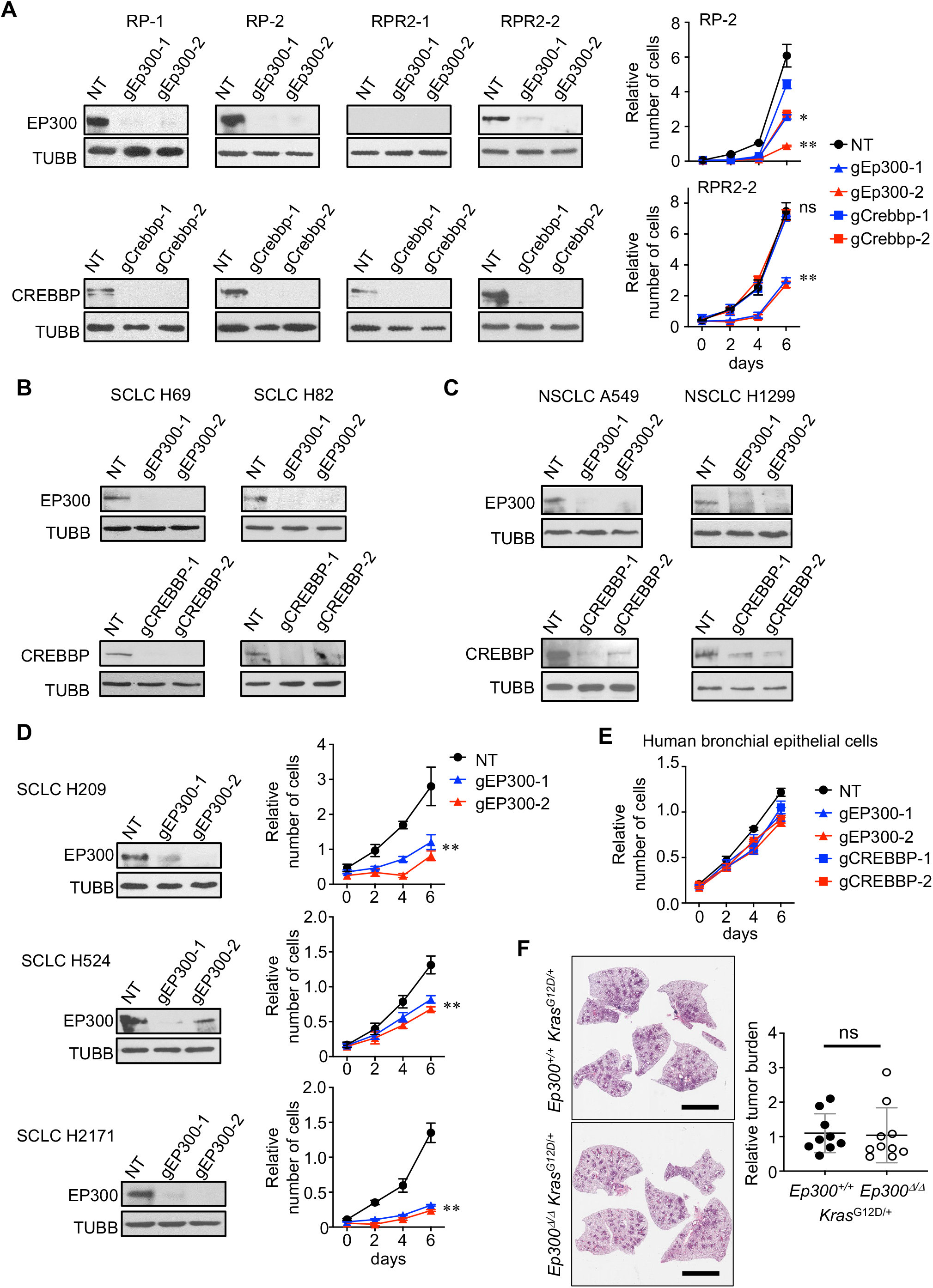
EP300 and CREBBP knockout in mouse SCLC cells and human cell lines, and effect of EP300/CREBBP knockouts on cells. **A-D,** Representative immunoblots for EP300 and/or CREBBP in mouse SCLC cells (A), human SCLC cell lines (B, D), and human NSCLC cell lines (C) infected with lentivirus expressing Cas9 and gRNAs against the target genes. RP-1 and RP-2 cells were derived from *Rb1/Trp53*-mutant tumors, and RPR2-1 and RPR-2 were derived from *Rb1/Trp53/Rbl2*-mutant tumors. TUBB blots was used as a protein loading control. **A, D, E,** Results of CellTiter-Glo assay for mouse and human SCLC cells (A, D) and human bronchial epithelial cells (E) infected with lentiviral vectors expressing CRISPR/Cas9 and the indicated guide RNAs. **F,** Representative images of H&E-stained lung sections of *Ep300*^+/+^ *vs. Ep300*^∆/∆^ *Kras^G12D^* mice (left) and quantification of tumor burden (tumor area/lung area) (n=9 and n=10 per genotype) (right). Statistical tests performed using unpaired t-test. Error bars represent standard deviation. NT: non-targeting guide RNA. (n=3 per cell type). **, p<0.05; **, p<0.01. Statistical tests performed using unpaired t-test (ns: not significant). Error bars represent standard deviation. Scale bars: F, 5 mm.

**Figure S3.**
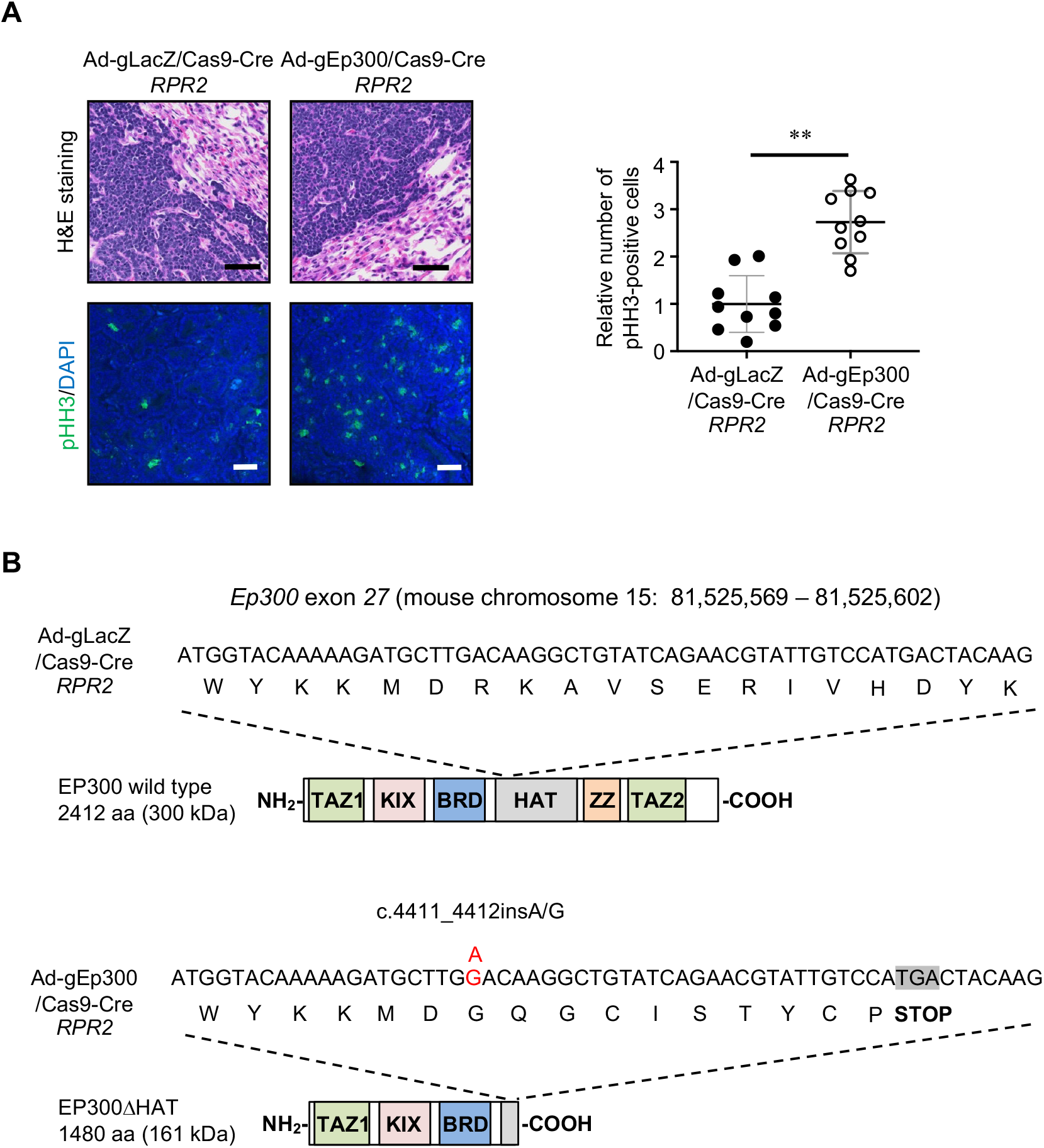
Targeted mutations in Ep300 exon 27 found in the tumors developed in mice. **A,** Representative images of H&E-stained lung sections of *RPR2* mice infected with Ad-gLacZ/Cas9-Cre *vs*. Ad-gEp300/Cas9-Cre virus (top), immunostaining for pHH3 (bottom), and quantification of pHH3-positive cells per tumor area (n=10 tumors per genotype) (right). DAPI (blue) stains for nuclei. **B**, Results of Sanger sequencing of the region in *Ep300* exon 27 targeted by the guide RNA, aligned with the diagram of wild-type and mutant EP300 proteins. Tumors developed in *RPR2* mice infected with Ad-gEp300/Cas9-Cre harbor a c.4411_4412insA/G mutation resulting in a premature stop codon. **, p<0.01. Statistical tests performed using unpaired t-test. Error bars represent standard deviation. Scale bars: A, 50 mm.

**Figure S4.**
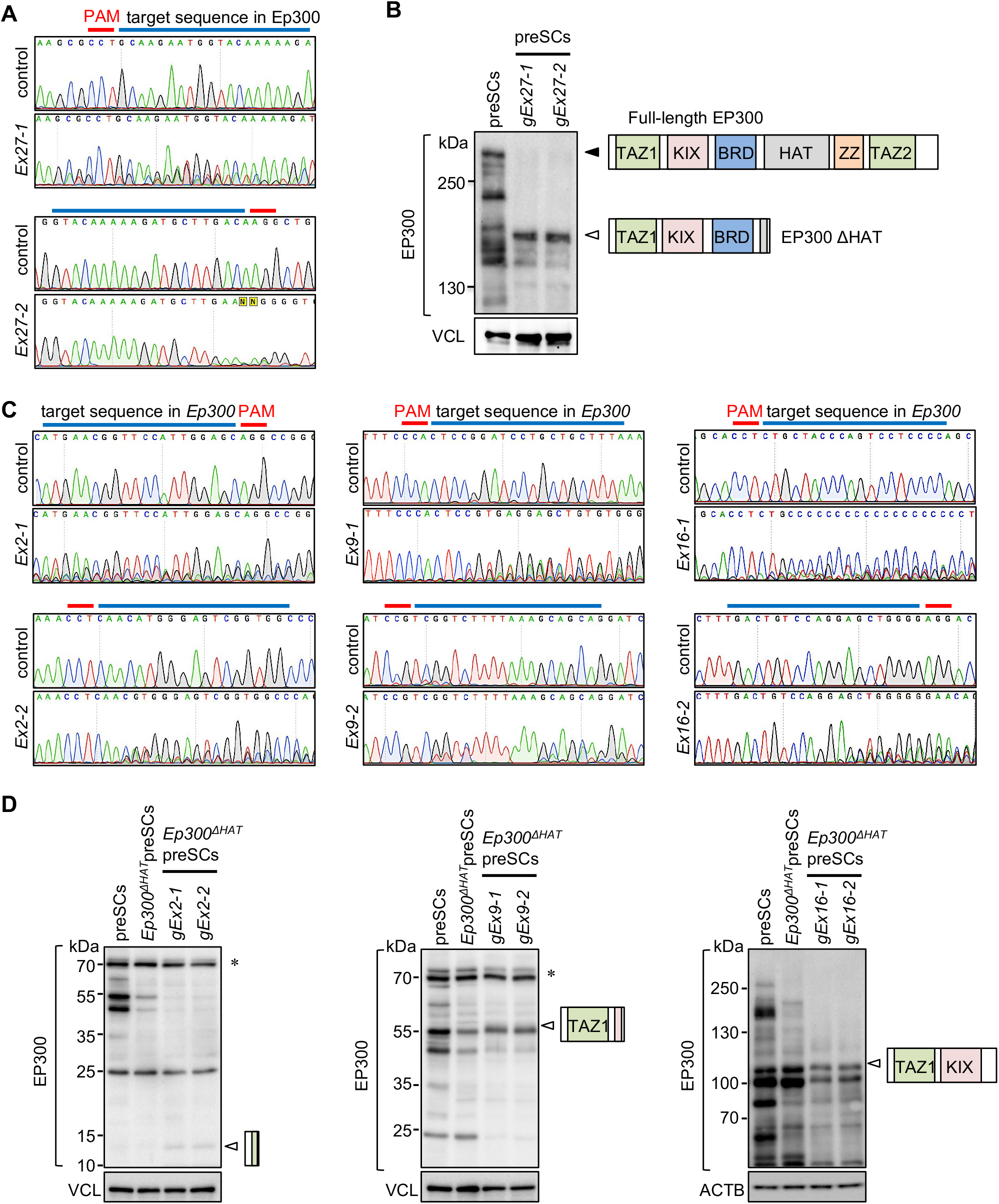
Validation of CRISPR/Cas9-mediated mutations in *Ep300* exons. **A, C,** Sequencing chromatograms showing the gRNA target sequences in control *vs. Ep300* exon 27-targeted preSCs (A) and *Ep300*^∆HAT^preSCs *vs.* Ep300 exons 2, 9, 16-targeted *Ep300*^∆HAT^preSCs (B). **B, D,** Immunoblot for EP300 in cells. Open arrowheads indicate bands of truncated EP300 variants that are generated as results of CRISPR-targeting. VCL and ACTB blots were used for a protein loading control. Asterisks indicate non-specific bands.

**Figure S5.**
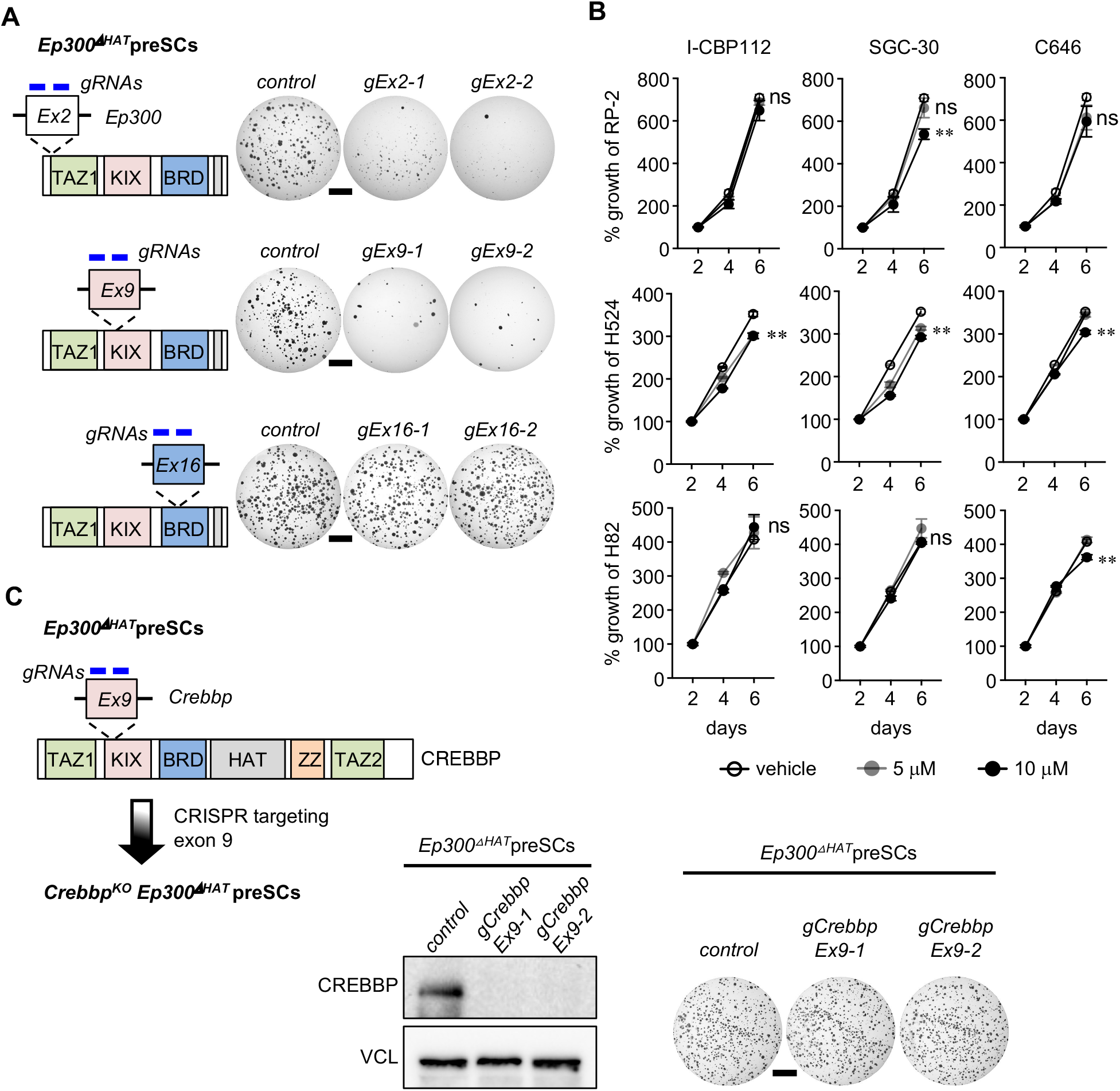
*Ep300*^∆HAT^preSCs and *Crebbp^KO^ Ep300*^∆HAT^preSCs and disrupting functions of EP300 and CREBBP KIX domains. **A** Schematics of CRISPR-mediated targeting of exons 2, 9, and 16 (left) and representative images of soft agar colonies formed from targeted cells (right). **B,** Results of MTT cell viability assay of mouse SCLC cells (RP-2) and human SCLC cell lines treated with selective bromodomain (BRD) inhibitors of CREBBP/EP300 (I-CBP112 and SGC-CBP30) and a selective histone acetyltransferase domain (HAT) inhibitor of CREBBP and EP300 (C646) (n=3 per cell type). **C,** Schematic of generating *Crebbp^KO^Ep300^∆HAT^*preSCs (left), Immunoblot for CREBBP in CRISPR-targeted *Ep300^∆HAT^*preSCs (middle), and representative images of soft agar colonies formed from CRISPR-targeted cells (right). VCL was used as a protein loading control. **, p<0.01. Statistical tests performed using unpaired t-test (ns: not significant). Error bars represent standard deviation. Scale bars; A, C, 5 mm.

**Figure S6.**
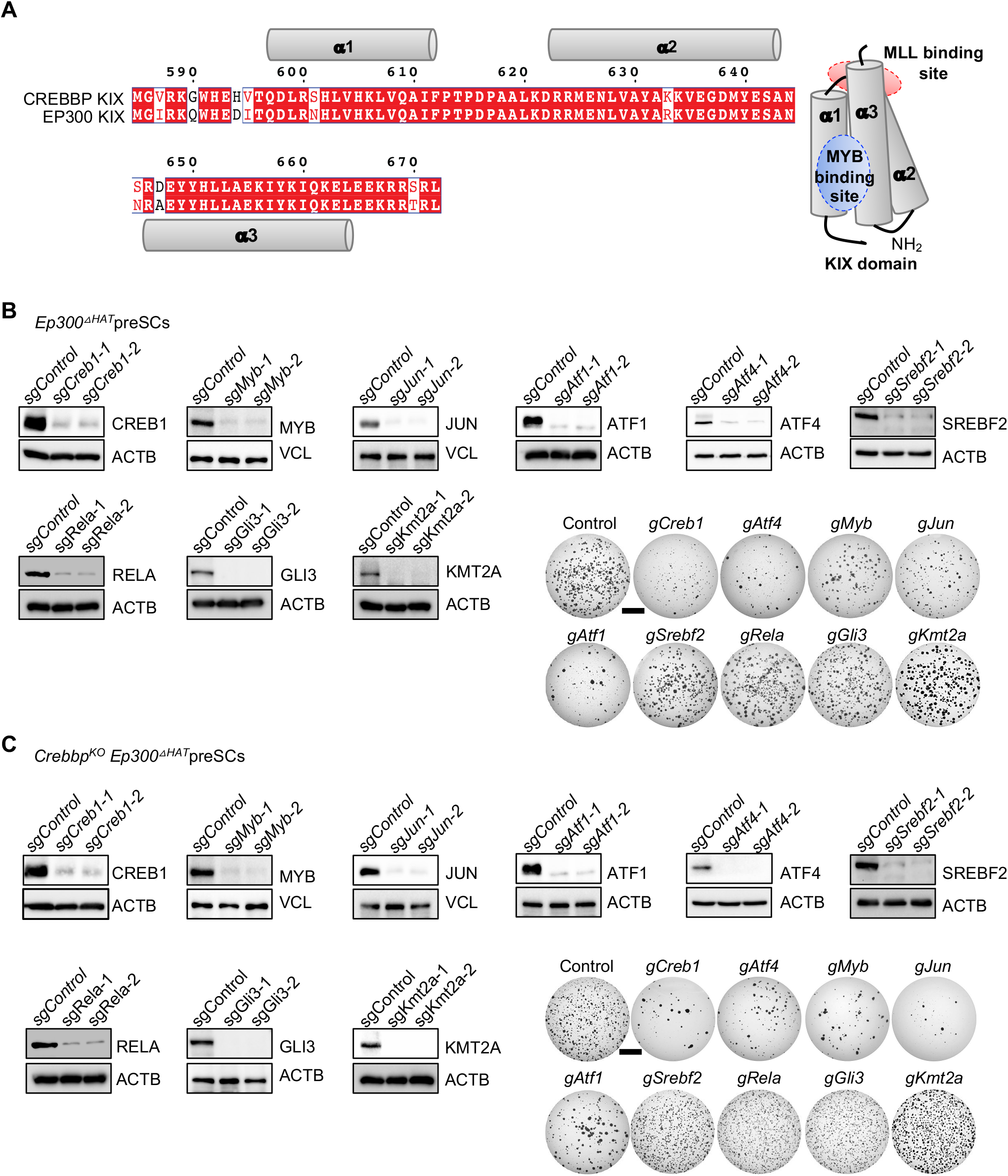
Structure of EP300/CREBBP KIX domains and knockout of transcription factors known to interact with the KIX domains of EP300 and CREBBP in *Ep300*^∆HAT^preSCs and *Crebbp^KO^ Ep300*^∆HAT^preSCs. **A,** Alignment of the amino acid sequences of human CREBBP and EP300 KIX domains. Numbering of residues is based on the CREBBP KIX domain (left). Cylinder representation of the structure of the KIX domain with MLL and MYB binding sites (right). **B, C.** Immunoblots validate loss of EP300/CREBBP KIX domain-interacting partner proteins (left) and representative images of soft agar colonies formed from targeted cells as indicated (right). ACTB and VCL blots were used as protein loading controls. Scale bars: B, C, 5 mm.

**Figure S7.**
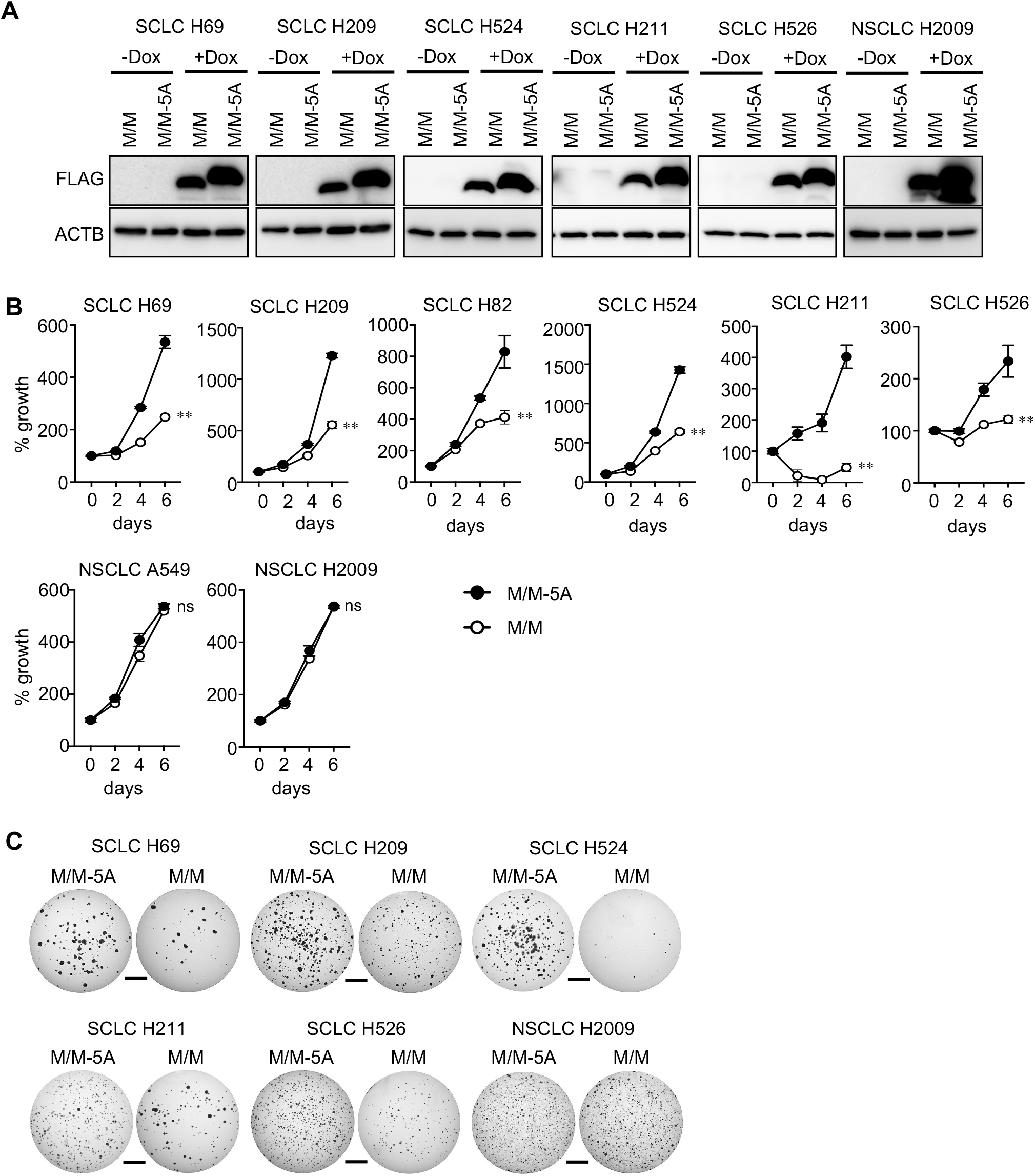
A peptide blocker of KIX-mediated protein interactions inhibits SCLC cell proliferation, but not NSCLC cells. **A,** Immunoblots for FLAG tag in M/M or M/M-5A peptides in SCLC and NSCLC cell lines expressing FLAG-tagged M/M or M/M-5A peptides under the control of a doxycycline (dox)-inducible promoter. ACTB blot was used as protein loading control. **B,** Results of MTT cell viability assay of human SCLC and NSCLC cell lines expressing M/M or M/M-5A (n=3 per cell type). **C,** Representative images of soft agar growth of human SCLC and NSCLC cell lines expressing M/M or M/M-5A (n=3 per cell type). **, p<0.01. Statistical tests performed using unpaired t-test. ns: not significant. Scale bars: C, 5mm.

**Figure S8.**
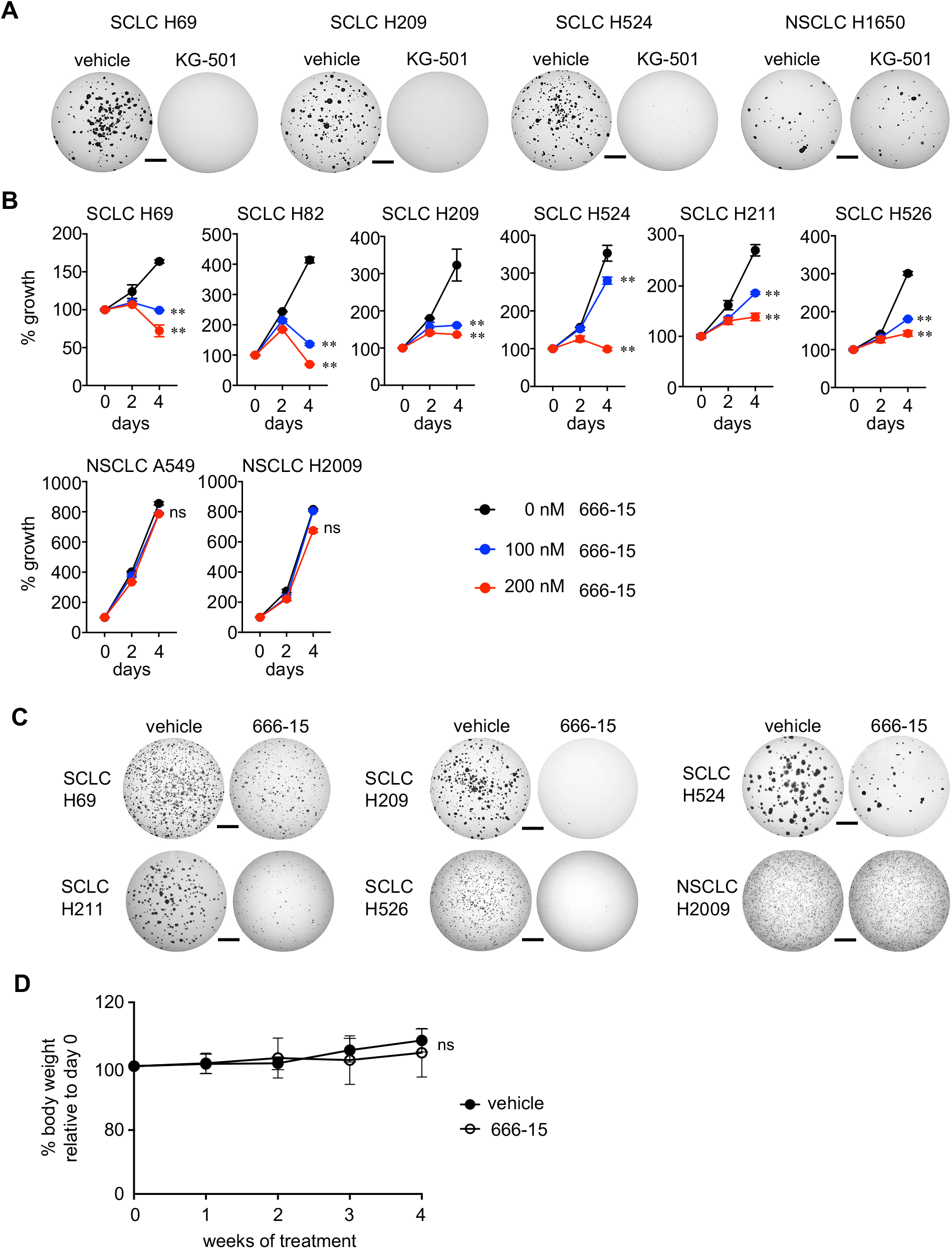
A chemical inhibitor of KIX-mediated protein interactions inhibits SCLC cell proliferation, but it does not inhibit proliferation of NSCLC cells and does not cause weight loss in mice. **A,** Representative images of soft agar growth of human SCLC and NSCLC cell lines treated with 2 mM of KG-501 or vehicle (n=3 per cell type). **B,** Results of MTT cell viability assay of human SCLC and NSCLC cell lines treated with 100 and 200 nM concentrations of 666-15 or vehicle (n=3 per cell type). **C,** Representative images of soft agar growth of human SCLC and NSCLC cell lines treated with 200 nM of 666-15 or vehicle (n=3 per cell type). **D,** Plot of % body weight of the mice injected with mSCLC RP-1 cells and treated with 666-15 or vehicle during 4-week period treatment. Daily treatment of 666-15 (10mg/kg) does not cause weight loss. **, p<0.01. Statistical tests performed using unpaired t-test. ns: not significant. Scale bars: A, C, 5mm.

**Figure S9.**
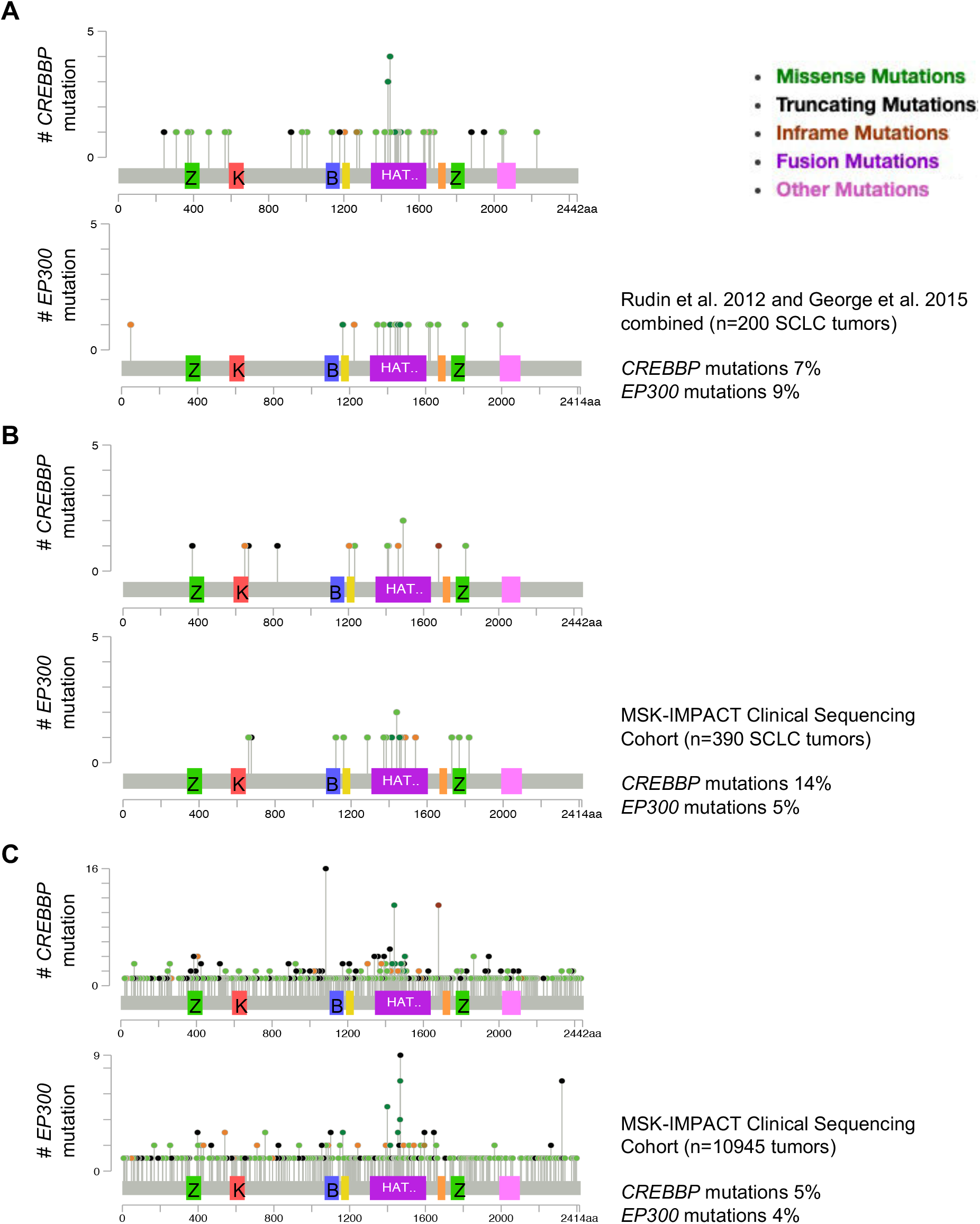
The patterns of EP300 and CREBBP mutations in human tumors. **A-C,** Lollipop diagrams generated from cBioportal show the distribution of mutations in *EP300* and *CREBBP* genes with respect to the protein structures. In both the published data from George et al. and Rudin et al. (references 7 and 9) (A) and the MSK-IMPACT clinical sequencing cohort (B), mutations that would result in loss of function are lacking in the coding sequences for the EP300 KIX domain but occur in the coding sequences for CREBBP KIX domain. In lymphoma and other cancer types, the mutations patterns in *EP300* and *CREBBP* are similar (C). Z: TAZ domain, K: KIX domain, B: Bromodomain, HAT: HAT domain.

