## Supplementary Information for "Unique role and vulnerability of EP300 KIX domain in small-cell lung cancer"

**Antibodies**

| Antibodies | Source, Product Number | Analysis, Antibody titer |
| --- | --- | --- |
| anti-CREBBP | Cell Signaling Technology, 73899 | Immunoblot, 1:2000 |
| | Thermo Fisher Scientific, PA1-847 | Immunoblot, 1:1000 |
| | Santa Cruz Biotechnology, sc-369 | Immunoblot, 1:2000 |
| anti-EP300 | Bethyl Laboratories, A300-358A | Immunoblot, 1:2000 |
| | GeneTex, GTX56157 | Immunoblot, 1:2000 |
| | Santa Cruz Biotechnology, sc-585 | Immunoblot, 1:2000 |
| anti-CREBBP | Cell Signaling Technology, 9104 | Immunoblot, 1:2000 |
| anti-RELA | Cell Signaling Technology, 8242 | Immunoblot, 1:2000 |
| anti-KMT2A (MLL) | Cell Signaling Technology, 14689 | Immunoblot, 1:1000 |
| anti-JUN | Cell Signaling Technology, 9165 | Immunoblot, 1:2000 |
| anti-SREBF2 (SREBP2) | Thermo Fisher Scientific, PA1-338 | Immunoblot, 1:2000 |
| anti-GLI3 | Thermo Fisher Scientific, PA5-28029 | Immunoblot, 1:1000 |
| anti-ATF4 | Santa Cruz Biotechnology, sc-200 | Immunoblot, 1:1000 |
| anti-ATF1 | Santa Cruz Biotechnology, sc-243 | Immunoblot, 1:1000 |
| anti-MYB | Santa Cruz Biotechnology, sc-74512 | Immunoblot, 1:1000 |
| anti-YY1 | Santa Cruz Biotechnology, sc-7341 | Immunoblot, 1:5000 |
| anti-ACTB | Santa Cruz Biotechnology, sc-47778 | Immunoblot, 1:5000 |
| anti-VCL (VINCULIN) | Santa Cruz Biotechnology, sc-73614 | Immunoblot, 1:5000 |
| Normal IgG | Santa Cruz Biotechnology, sc-2025 | |
| anti-TUBB | Sigma, T8328 | Immunoblot, 1:2000 |
| anti-FLAG | Sigma, F1804 | Immunoblot, 1:10000 |
| anti-CALCA (CGRP) | Sigma, C8198 | Immunostaining, 1:100 |
| anti-phosphorylated histone H3 (pHH3) | Abcam, Ab1791 | Immunostaining, 1:100 |
| Secondary Ab-488 | Thermo Fisher Scientific, A11034 | Immunoblot, 1:200 |
| Scondary Ab-HRP (mouse) | Jackson Immuno Research, 115-035-003 | Immunoblot, 1:5000 |
| Secondary Ab-HRP (Rabbit) | Jackson Immuno Research, 111-035-003 | Immunoblot, 1:5000 |

**Plasmids**

| Plasmid | Source | Selection markers |
| --- | --- | --- |
| pCW57.1 | Addgene, 41393 | Puromycin |
| psPAX2 | Addgene, 12259 | not applicable |
| pMD2.G | Addgene, 12260 | not applicable |
| pCW-Cas9 | Addgene, 50661 | Puromycin |
| pL-CRISPR.EFS.tRFP | Addgene, 57819 | not applicable |
| LV-gRNA-zeocin | gift from Mazhar Adli | Zeocin |
| pMAL-c5E | NEB, N8110 | not applicable |
| pET32a | GenScript | not applicable |
| pCW57.1-3XFLAG-M/M | this study | Puromycin |
| pCW57.1-3XFLAG-M/M-5A | this study | Puromycin |

**Guide RNAs**

| Target gene (exon number) | Species | Sequence (5' to 3') |
| --- | --- | --- |
| Non target | not applicable | GGGATACTTCTTCGAACGTTT |
| *CREBBP* (exon 2) | Human | CGCGGGACTGAACACCGCAC |
| | | GCAGCCGAACAGTGCTAACA |
| *EP300* (exon2 ) | Human | GTTCAATTGGAGCAGGCCGA |
| | | GAATTGGGACTAACCAATGG |
| *Crebbp* (exon 2) | Mouse | GGCCTTCTCAATAGTAACTC |
| | | AGCGGCTCTAGCATCAACCC |

| | | |
|---|---|---|
| *Ep300* (exon 24) | Mouse | GATTGCCATCTACCAGACTT |
| *Ep300* (exon 27) | Mouse | GTACAAAAAGATGCTTGACA |
| *Ep300* (exon 2) | Mouse | ATGAACGGTTCCATTGGAGC |
| | | GGCCACCGACTCCCATGTTG |
| *Ep300* (exon 9) | Mouse | TAAAGCAGCAGGATCCGGAG |
| | | TGTTGCATATGCTCGTAAAG |
| *Ep300* (exon 16) | Mouse | TGGGGAGGACTGGGTAGCAG |
| | | TGACTGTCCAGGAGCTGGGG |
| *Creb1* (exon 2) | Mouse | CAGCTGCACTAAGGTTACAG |
| | | GACCTGGACTGTCTGCCCAT |
| *Atf4* (exon 1) | Mouse | CGAGGAGCCCGCCTTGTCGC |
| | | GAACAGCGAAGTGTTGGCGG |
| *Myb* (exon 2) | Mouse | AATCTGGAAAGCGTCACTTG |
| | | TGTGTGACCATGACTACGAT |
| *Jun* (exon 1) | Mouse | CTTCTCACGTCGCCCGACGT |
| | | CGAGCAGGAGGGCTTCGCCG |
| *Elk4* (exon 1) | Mouse | AGGAGGTGGCTCGTCTTTGG |
| | | GTTCCTTCTCCAGCTCCTGC |
| *Atf1* (exon 1) | Mouse | TCTCTGTCGTGTTACTCTTG |
| | | AGCCTGGTTCAACCGTTGCG |
| *Srebf2* (exon 2) | Mouse | AGTGGCAGAGGCAACAATGG |
| | | CCACAGACCCTGCAGTACAG |
| *Yy1* (exon 1) | Mouse | GATGTAGAGGGTGTCGCCCG |
| | | ACGCGCGAGGAGGTGGTCGG |
| *Rela* (exon 3) | Mouse | TAAATGCGAGGGGCGCTCAG |
| | | CCAGGCTTCTGGGCCTTATG |
| *Gli3* (exon 4) | Mouse | ATACGTCGGGCTACTAGATA |
| | | GGGGTGTGGGGGGCTGAAGG |
| *Kmt2a* (exon35) | Mouse | CAGTTCCTTCTGACCGCCAC |
| | | TCAAAGCTTAGCATGATCCT |
| *Kmt2a* (exon 2) | Mouse | TTTCATCTAGGCCTTTCCTT |
| | | AGCCCCACCAGGTCTCCTTC |

**Guide RNAs used for Ad-CRISPR-Cre vectors**

| Target gene | Species | Sequence (5' to 3') |
|---|---|---|
| *LacZ* | E. coli | GGCTGCGCAACTGTTGGGAA |
| *Ep300* (exon 27) | Mouse | GTACAAAAAGATGCTTGACA |

**Sequencing primers**

| Target gene | Target exon | Sequence (5' to 3') |
|---|---|---|
| *Ep300* | exon 2 (forward) | GGTTCACTGTTTGACCTGGAA |
| | exon 2 (reverse) | AGGAGAGCCCTGCTGTAGTG |
| | exon 9 (forward) | CTTCAAATTAAGTGTGGGGTTTTT |
| | exon 9 (reverse) | AGCACCTGGGAGTCAGAGAC |
| | exon 16 (forward) | TTCCTGTCTTGCCTTTCAGG |
| | exon 16 (reverse) | TCCCAAGCACTGGGATTAAA |
| | exon 27 (forward) | CTACTCTAGGAGGACTACTGTAGCG |
| | exon 27 (reverse) | GTCTACAAAGTGAGTTCCAGGGTAG |

**Genotyping primers**

| Target alleles | | Sequence (5' to 3') |
|---|---|---|
| *Trp53 lox/lox* | forward | CACAAAAACAGGTTAAACCCAG |
| | reverse | AGCACATAGGAGGCAGAGAC |
| *Rb1 lox/lox* | forward | CTCTAGATCCTCTCATTCTTCCC |
| | reverse | CCTTGACCATAGCCCAGCAC |
| *Rbl2 lox/lox* | forward | GTGTTGTAACATTCTCGTGGG |
| | reverse | GACTGCTGGTATTAGAACCC |
| *Ep300 lox/lox* | forward | GTGAGTTGATGTCCCTGTCG |
| | reverse | CAGACACCCTCTTGCACTCA |
| *H11 lox-STOP-lox-Myc T58A* | MycT58A-forward | AACTTCCCGCCGCCGTTGTT |
| | MycT58A-reverse | CAACGGGCCACAACTCCTCA |
| | H11-forward | TGGAGGAGGACAAACTGGTCAC |
| | H11-reverse | TTCCCTTTCTGCTTCATCTTGC |
| *Kras lox-STOP-lox-G12D* | wild type forward | TGTCTTTCCCCAGCACAGT |
| | mutant forward | GCAGGTCGAGGGACCTAATA |
| | common reverse | CTGCATAGTACGCTATACCCTGT |
| *Rosa26 lox-STOP-lox-luciferase* | wild type forward | CGTGATCTGCAACTCCAGTC |
| | wild type reverse | GGAGCGGGAGAAATGGATATG |
| | mutant forward | CCAGGGATTTCAGTCGATGT |
| | mutant reverse | AATCTGACGCAGGCAGTTCT |
